# TopoMIL: Topology Improves Multiple Instance Learning in Diagnostic Microscopic Images

**DOI:** 10.64898/2026.06.10.731443

**Authors:** Salome Kazeminia, Muhammed Furkan Dasdelen, Bastian Rieck, Carsten Marr

## Abstract

Microscopic images of cells and tissues are central to disease diagnosis. In computational pathology, multiple instance learning (MIL) has emerged as a key paradigm for analyzing numerous images within a single patient sample. While the representative distribution of cells in a sample is important for diagnosis, existing MIL frameworks largely overlook it. We introduce TopoMIL, a framework that extracts the representative topological structure of the sample and integrates it into the MIL classifier. Three topological representations are assessed, each with distinct advantages and computational costs. We evaluate TopoMIL on four histopathology and cytomorphology datasets, each presenting unique challenges. Integrating the sample’s topological information into MIL enhances classification across average, max, attention-based, and transformer pooling, yielding AUCROC gains of 3.3%, 4.2%, 5.9%, and 0.5%, respectively, with moderate computational cost. Our work underscores the potential of TopoMIL as a scalable extension to existing morphology-based models in computational pathology.

## Introduction

The analysis of high-resolution microscopic images of cells and tissues is critical for the identification and diagnosis of cancer^1–4^. However, these images may encompass thousands of microscopic regions, the vast majority of which exhibit normal tissue and cells, while a minority contain pathological alterations that are relevant for diagnosis. This challenge has motivated the use of machine learning methods capable of efficiently identifying and analyzing diagnostically relevant information within the samples. To enable the application of machine learning techniques, these images are partitioned into smaller subregions, or “instances,” which together constitute a “bag” with a single label reflecting the diagnostic outcome assigned to the entire high-resolution image. Multiple Instance Learning (MIL) is a widely adopted machine learning paradigm well-suited for this type of weakly labeled data^5,6^. MIL frameworks usually consist of three key components: (i) an instance encoder, which extracts morphological features from individual instance images, (ii) an aggregator, to combine instance-level features and generate a bag-level morphological representation, and (iii) a classifier head to predict the label of the whole bag (Fig. 1a).

**Figure 1:**
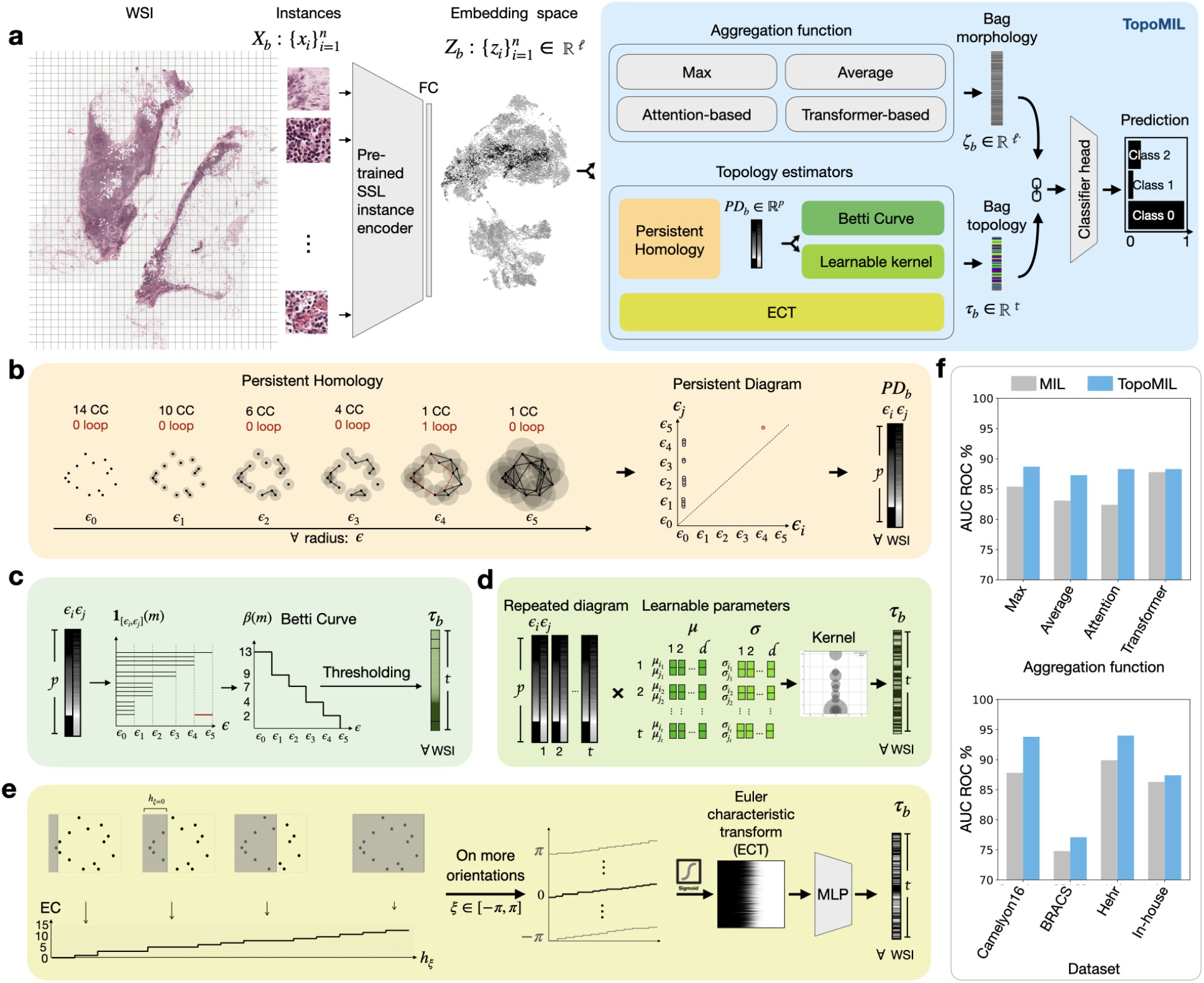
TopoMIL integrates topology into multiple instance learning. **a** A pretrained instance encoder maps each image *x*_*i*_ within a bag *X*_*b*_ to a latent feature space *Z*_*b*_. Anaggregator then pools the instance-level features into a unified morphological representation ζ_*b*_ in *l*’-dimensional space (*l*’ 384 for cytomorphology and 1024 for histopathology data, based on the foundation feature extractor). In parallel, the topology estimation block computes a topological representation of the bag τ_*b*_ in *t* dimensional space. The concatenated feature vector is fed into the classifier, which, e.g., predicts a patient’s disease. **b** Persistent homology (PH) computation tracks the emergence and disappearance of topological features by expanding the neighborhood distance around instance data points, yielding persistence diagrams *PD*_*b*_ with birth–death scale pairs (ϵ_*i*_, ϵ_*j*_ ). *PD*_*b*_ needs to be represented as a vector to be used in the model. **c** Betti curve representation captures the evolution of topological features over fixed distance thresholds derived from the *PD*_*b*_ . **d** Topological representation kernel learns the representation of *PD*_*b*_ probabilistically, encoding it as a vector of learned mean (µ) and standard deviation (σ) parameters. **e** The Euler Characteristic Transform (ECT) captures the topological structure of the bag’s point cloud by scanning the latent space across multiple directions. The resulting 2D ECT is embedded into a feature vector using a multi-layer perceptron (MLP). **f** Averaged performance of TopoMIL (the best performance) compared to MIL.

Recent advances in foundation models have contributed to a rich extraction of morphological features within the instance encoders of MIL frameworks^7,8^. In parallel, advances in MIL largely concentrated on developing more effective aggregation functions, such as attention-based pooling^9^ and transformer-based architectures^10^, to identify and emphasize the most class-relevant morphological features within a bag. Despite the ability of transformers to model some relational information between instances, the explicit estimation of topological structure of bags remains generally overlooked across MIL research, while such information is recognized as clinically important^11–16^.

Modeling the topology of instances within a bag in MIL is challenging, particularly in the absence of instance-level annotations and given the intricate arrangement of the underlying feature space. In histopathology, instances (image patches) each encompass a collection of cells that may be homogeneous, composed of a single cell type, or heterogeneous, containing mixtures of morphologically and functionally distinct cells. This variability in intra-patch composition can lead to embeddings that are not clearly separable, particularly when multiple cell types co-occur within a patch, and may introduce substantial noise or obscure diagnostically relevant information within the patch. In cytomorphology, where instances represent individual white blood cells^17^, the feature space often reflects the arrangement of biological variation. Optimal embeddings lie along continuous low-dimensional manifolds associated with hematopoietic differentiation trajectories, while mature cell types form discrete clusters corresponding to terminal phenotypes^8^. Characterizing the structural arrangement of instances is complicated by inherent noise and variability in the data, some of which may not be fully addressed by feature extractors. In addition, batch effects, domain shifts, and the presence of rare or previously unseen instance types can distort the feature space, particularly in clinical settings^18^.

Topological data analysis (TDA) characterizes the global shape and arrangement of data point clouds, without relying on parametric assumptions, such as predefined clustered or specific distributional forms^19^. Moreover, TDA’s robustness to noise and perturbations, combined with its flexibility in capturing structural variability, enables it, within certain limits, to address variability and noise inherent in the data that are not fully mitigated by feature extractors^20,21^. While TDA has been applied in MIL for cytomorphology and histopathology data, previous studies have mainly focused on extracting topological features from instance-level images, representing morphological structures as pixel-based point clouds^22,23^, rather than capturing the topological structure of the entire bag based on instance embeddings. The only study addressing bag-level topological structure derived from instance representation spaces employs topological regularization to align data and feature spaces, thereby improving instance embeddings under simple and scarce data scenarios, without leveraging bag-level topological structure for classification^24^.

We introduce TopoMIL, a topology estimation module that characterizes the arrangement of instances within each bag and utilizes it for classification. We consider three different representations of extracted topological features: (i) Persistent homology (PH) with Betti curve summaries, (ii) PH with a learnable representation kernel, and (iii) Euler Characteristic Transformation (ECT). The first two approaches employ PH to extract topological features from the instance arrangement, resulting in a persistence diagram (Fig. 1b). However, they differ in their vector representations: the Betti curve (Fig. 1c) generates a statistical embedding using fixed functions and thresholds, while the learnable kernel representation (Fig. 1d) enables the modeling of the probability distribution of topological features tailored to the classification objective. PH, while tolerant to noise and small perturbations in the data, is computationally complex; therefore, we also consider the Euler characteristic transformation as a less demanding alternative for representing the topological structure of instance arrangement. This approach summarizes the global shape of the instance arrangement by counting data points along multiple directions in the feature space (Fig. 1e). While it does not explicitly capture topological features and is less robust to noise and deformations (compared to PH), it offers a differentiable detailed shape representation with significantly lower computational cost.

We evaluate the effectiveness of these three approaches on two histopathology and two cytomorphology classification tasks, which have different characteristics and challenges. Averaged over all topological descriptors and aggregation functions, TopoMIL improves MIL performance by 6% and 2.3% on the histopathology datasets and by 4.1% and 1.1% on the cytomorphology datasets (Fig. 1f).

Our framework is designed modularly, enabling seamless integration of topological formulations into MIL architectures regardless of their aggregation function. Our results demonstrate that TopoMIL enhances the MIL model’s ability to exploit structural patterns in feature space, thereby improving classification.

## Results

We benchmark TopoMIL against conventional MIL models using different aggregation functions on four clinically relevant tasks: two in histopathology and two in cytomorphology.

### From solid tissue to peripheral blood

Histopathology datasets **Camelyon16**^25^ (Fig. 2a) and **BRACS**^26^ (Fig. 2b) consist of whole-slide images (WSIs) of solid tissue, characterized by densely packed cellular structures. Unlike Camelyon16 (introducing a binary classification task of normal vs. tumor WSI), where tumors typically appear as focal regions within largely normal tissue, BRACS exhibits high morphological diversity and substantial intra-class heterogeneity (among 3 classes). Since only small discriminative regions within each slide are diagnostically informative, BRACS represents one of the most challenging datasets for automated classification^26^.

**Figure 2:**
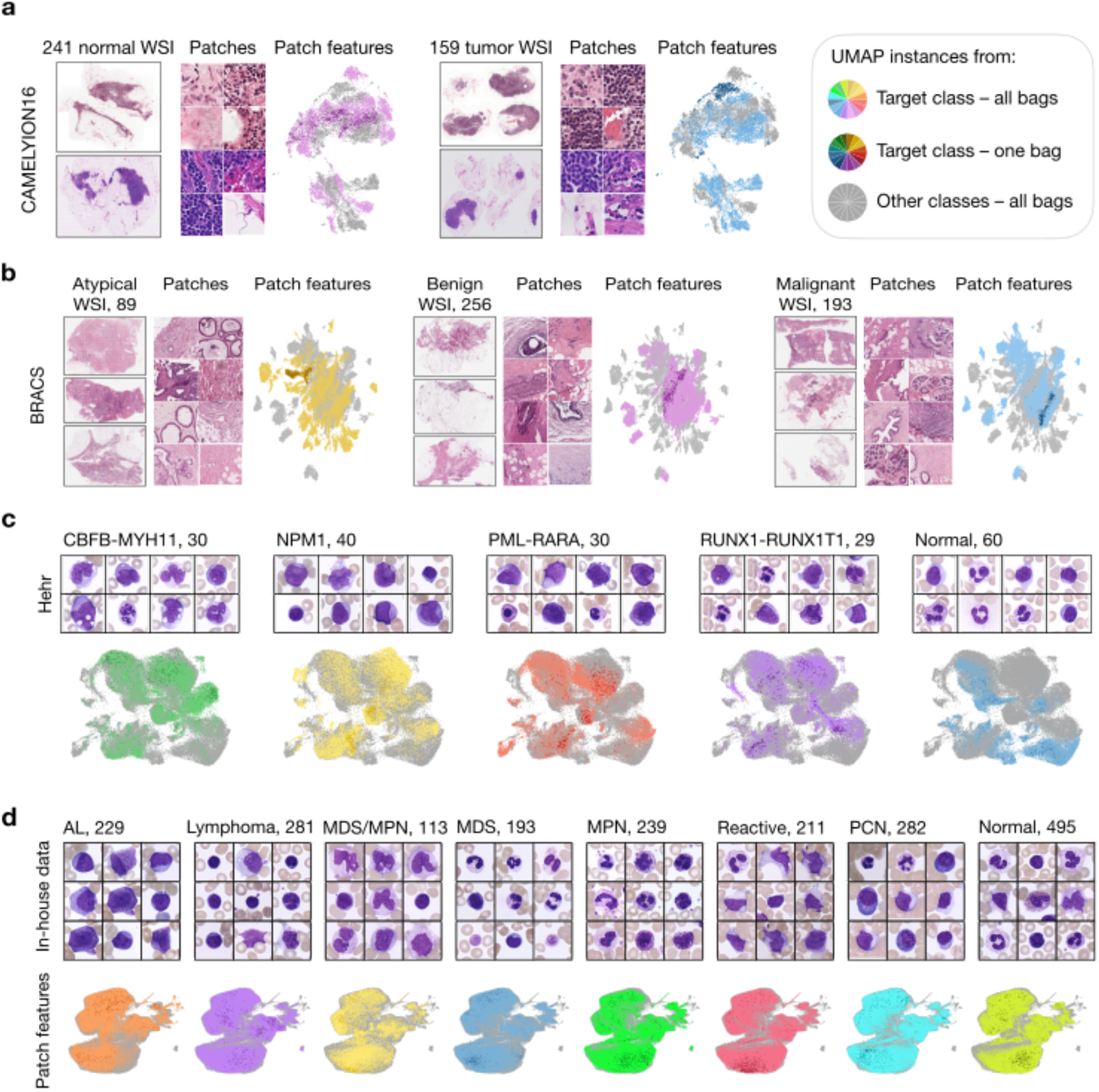
We evaluate TopoMIL on two histology datasets and two cytomorphology datasets. **a** CAMELYON16: 400 H&E-stained WSIs of lymph node sections (241 normal, 159 tumor). Shown are a representative WSI, example patches, and a 2D UMAP of UNI-extracted features. **b** BRACS: 547 WSIs from breast tissue biopsies (89 atypical, 265 benign, 193 malignant), with a representative WSI, image patches, and UMAP of UNI-extracted features. **c** Hehr: 189 digitized stained blood smears spanning five diagnostic classes (CBFB-MYH11: 30, NPM1: 40, PML-RARA: 30, RUNX1-RUNX1T1: 29, normal: 60). Shown are example smear images and a 2D UMAP of DinoBloom-extracted cell-level features. **d** In-house data: peripheral blood single cell images collected from 2043 patients from 8 diagnostic classes. In all UMAPs, instances are colored by class, with one random sample per class highlighted in a darker shade.

We also apply our methods to the publicly available **Hehr** WBC cytomorphology dataset^27^ (Fig. 2c), as well as our **in-house** dataset^28^ (Fig. 2d). Challenges of Hehr dataset are semantic ambiguity due to the overlapping morphologies of class-relevant cells across five different genetic subtypes, high intra-class variability within the same genetic subtype, and the presence of atypical cells in some healthy samples resulting from non-malignant reactive conditions^27^. Notably, the training split of the Hehr dataset is used as one of the training datasets for DinoBloom^8^, underscoring the high quality of instance embeddings achieved in this setting. Our in-house dataset contains eight diagnostic categories that represent broad clinical groupings but are internally diverse, with substantial variability in subtypes and overlapping features across classes. Collectively, the high intra-class heterogeneity and inter-class overlap make this dataset particularly challenging for a robust classification. More explanation of the datasets is provided in the Supplementary Information Dataset.

### TopoMIL outperforms MIL across datasets and pooling strategies

Accurate classification of cellular images and image patches relies not only on image features, but also on the representetive topological structure of a sample. To test whether bag-level topological features can capture this complementary information, we compare the baseline MIL with TopoMIL, which integrates three topology-based descriptors (PH with Betti curves, PH with kernels, and ECT) across four aggregation strategies (average pooling, max pooling, attention-based pooling, and transformer-based pooling). Incorporating topological descriptors consistently enhances MIL performance across datasets and aggregation strategies (see Supplementary Tables 1, 2, and Methods for details).

To apply TopoMIL to the Camelyon16 dataset, we randomly sample 1000 instances per bag to keep topological computation feasible. The sampled subset is used to estimate the bag’s topological signature, serving as a representative approximation of its overall topological structure. Our experiments (Fig. 3a) show that max pooling yields the highest performance (97.7±1.0% AUROC) on the baseline MIL, suggesting that the most prominent cellular morphological feature is sufficient for effective classification in this dataset. This is followed by transformer-based pooling (92.1 ± 4.9% AUROC), average pooling (83.0 ± 6.6% AUROC), and attention-based pooling (78.2 ± 5.3% AUROC). For max pooling, TopoMIL improves the performance relative to the baseline by 2%, but the enhancement is not statistically significant.

**Figure 3:**
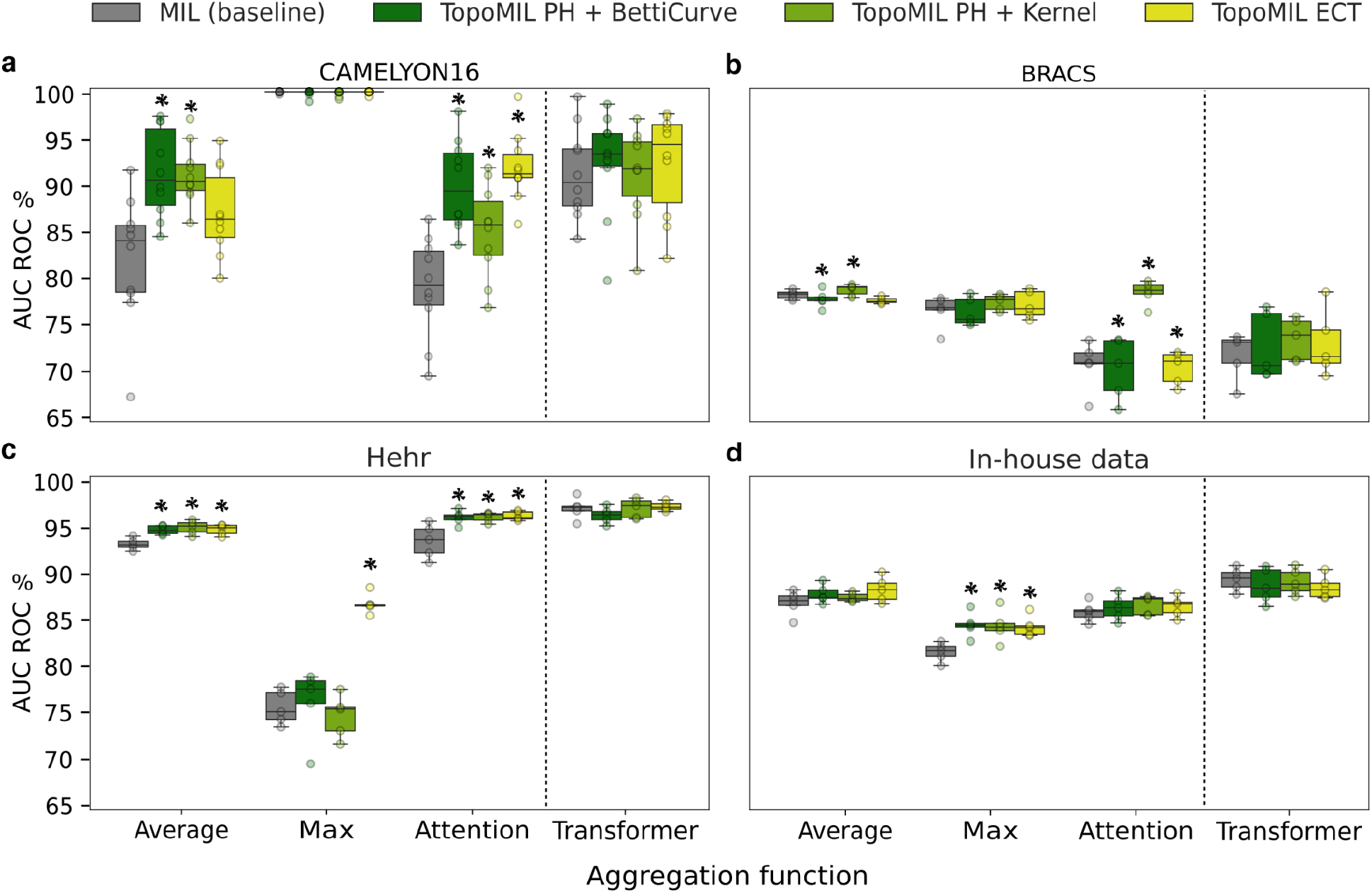
TopoMIL enhances MIL performance across four diverse biomedical imaging datasets. AUC-ROC for the Camelyon16 (**a**) and BRACS histopathology (**b**) datasets, and the Hehr (**c**) and our in-house (**d**) cytomorphology datasets. Each box-and-whisker plot shows the performance of the baseline MIL model and the TopoMIL framework using three topology embedding techniques: persistent homology (PH) with Betti curve representation, PH with learnable kernel representation, and Euler Characteristic Transform (ECT). Models are evaluated using four instance aggregation strategies: average pooling, max pooling, attention-based pooling, and transformer-based aggregation. Across all datasets and aggregation methods, incorporating topological information improves classification performance, demonstrating the benefits of topological representations in both histopathology and cytomorphology imaging domains. A statistically significant enhancement, as calculated by the Kruskal-Wallis test and compared to the baseline, is specified by *.

For average pooling, BettiCurve and kernel-based representations significantly improve performance relative to the baseline, with 9.8% (p-value = 0.0041 and 0.0011, Kruskal-Wallis test), while ECT provides a smaller improvement of 6.0%. In contrast, under attention pooling, all three TopoMIL approaches significantly enhance performance, with ECT showing the largest improvement (16.7%, p-value = 0.0002), followed by BettiCurve (15.6%, p-value = 0.0005) and the kernel-based representation (9.0%, p = 0.0282). Notably, the combination of attention pooling with topological representation outperforms TopoMIL with average pooling and achieves performance comparable to that of transformer-based pooling. To make transformer-based aggregation computationally manageable within TopoMIL, we subsampled 1% of instances per bag to form input bags. This baseline transformer still outperforms attention pooling by more effectively capturing relationships among instances. The inclusion of topological information into transformer-based pooling provides only minimal additional benefit under this setting: BettiCurve and ECT yield modest improvements of approximately 0.7%.

Our experiments on the BRACS dataset (Fig. 3b) show that average pooling yields the highest performance (78.3 ± 0.4% AUROC) on the baseline MIL, followed by max pooling (76.8 ± 1.6% AUROC), transformer-based pooling (72.5 ± 2.3% AUROC), and attention-based pooling (71.4 ± 2.5% AUROC). This indicates the importance of identifying and aggregating multiple class-relevant instances. Although attention mechanisms and transformer-based aggregation theoretically refine representations through learnable weighting, their underperformance in this context is probably due to the high inter-class diversity and the complexity of optimizing the topological representations of Betti curves and detailed representations of ECT. In contrast, the learnable kernel representation is capable of filtering out the irrelevant topological variations caused by intra-class heterogeneity. By preserving the robust class-relevant topological structure, the learnable kernel improves the performance of baseline MIL in all aggregations up to 8.0%.

On the cytomorphology Hehr dataset (Fig. 3c), transformer-based pooling works best (97.3±1.2% AUROC), followed by average pooling (93.1±0.6% AUROC) and attention-based pooling (93.8±1.6% AUROC). The lowest performance is achieved by max pooling (75.4±1.6% AUROC) that relies solely on the most prominent morphological features, which appears to be insufficient here. The strong instance representations provided by DinoBloom reduce the noise in the feature space, enabling ECT to improve baseline MIL significantly (2.1% to 14.9%, p=0.009 to 0.016). Importantly, TopoMIL improves MIL performance significantly for average pooling (p=0.009, 0.016 for PH with BettiCurve, Kernel respectively) and attention pooling (p=0.016, 0.016). This highlights the effectiveness of fine-grained topological information when anchored by key morphological features. In contrast, higher-level topological descriptors derived from PH, while still significantly enhancing average and attention-based pooling, yield comparatively smaller improvements of 2.4% and 4% over the baselines.

On the cytomorphology in-house dataset (Fig. 3d), the small gap between pooling strategies reflects both the less discriminative embedding space of instances and the overall difficulty of the classification task. From best to worst, transformer-based pooling achieves the highest performance (89.3±1.1% AUROC), followed by average pooling (87.2±1.3%), attention-based pooling (86.2±1.0%), and max pooling (82.4±1.0%). TopoMIL significantly improves MIL performance for max pooling (∼2.9%, p = 0.09, 0.03, 0.009 for PH with BettiCurve, PH with kernel, and ECT, respectively). Since topological features are estimated in the latent instance space, where each point’s position is defined by the instance encoder, the quality of the representation directly affects their usefulness. In this dataset, the less discriminative latent space limits the ability of topological representations to capture class-relevant patterns.

### Topology vs. Morphology

To quantify the contribution of the topological representation to the classification task, we perform an ablation analysis in which the morphological aggregation module is removed from the framework. In this configuration, the classifier predicts the bag label exclusively from the topological features of the bag. For this experiment, since the aggregation function is removed from the TopoMIL architecture, the reference performance is defined as that of the baseline MIL performance among all the tested aggregation strategies (see Fig. 4, with details in Supplementary information, Table 3).

**Fig 4:**
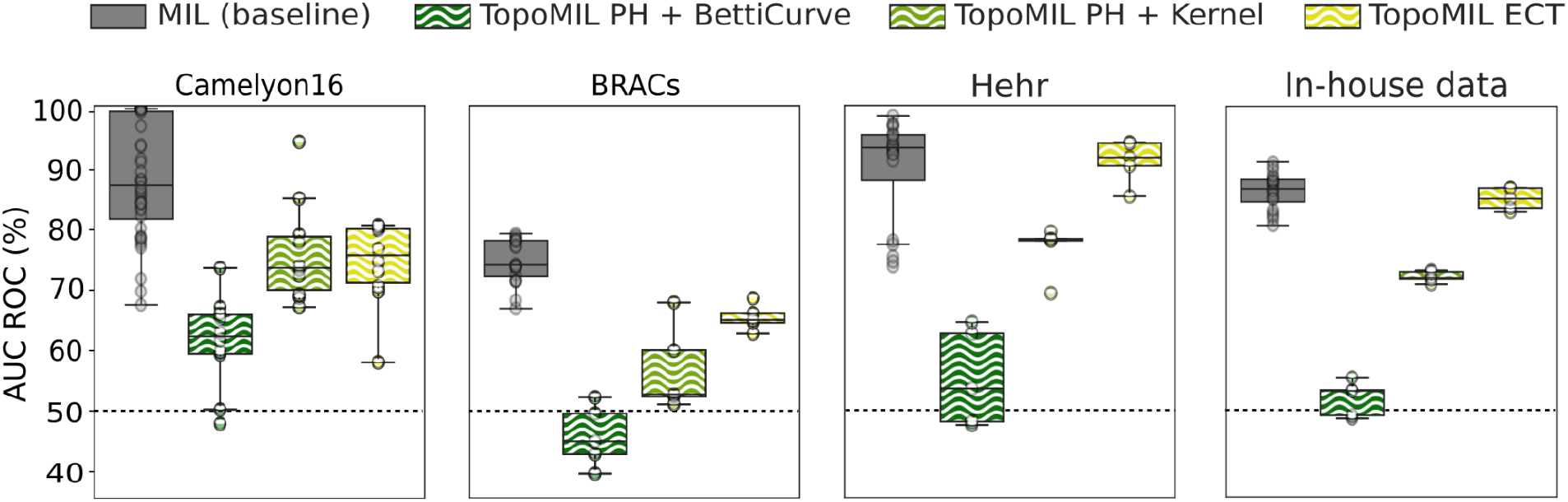
Ablation study illustrating the discriminative power of topological representation in MIL. TopoMIL (without morphological aggregation) relies solely on topological bag descriptors, while the baseline MIL models include only morphological aggregation. The MIL performance covers all four aggregation functions. The dashed line marks random performance.

Among the three approaches, PH, represented by the Betti curve, makes the lowest contribution and predicts close to random. We conclude that the fixed statistical calculation followed by fixed thresholding of the Betti curve does not adapt to the classification task and does not allow gradients to be propagated back to the instance representations. In contrast, PH, represented by the learnable kernel, provides flexibility to adapt to the data and the classification objective, which results in significantly higher performance. The ECT approach, capturing more detailed topological representation while remaining differentiable, also provides such flexibility and leads to the highest performance. Interestingly, on cytomorphology datasets, ECT provides comparable performance with the baseline MIL.

Using topology alone does not surpass baseline MIL. Topological features describe the bag’s overall shape in latent space by capturing instance arrangement and connectivity. Their invariance to translation, rotation, and scaling increases robustness but removes information about the bag’s absolute position. Consequently, bags with identical shapes, but different locations, may appear indistinguishable. Morphological features provide this positional context, anchoring the bag in latent space. Combining morphology and topology, therefore, integrates shape and context, yielding higher performance than either alone.

### Complexity and scalability

For TopoMIL, we use only 0-dimensional PH features. Their computational cost is dominated by pairwise distance computations between instances in a d-dimensional space, resulting in a time complexity of 𝒪 (*n*^2^ *d*) and memory complexity of 𝒪 (*n*^2^). The Betti curve and kernel-based variants add only minor overheads relative to PH. For the ECT, the time complexity is 𝒪 (*n*.*d*^2^) and the memory complexity of 𝒪 (*n* + *d*^2^). Pooling operations scale from linear (max/average) to quadratic (transformer), making transformer-based aggregation the most resource-demanding architecture. To assess the trade-off between performance and computational complexity, we measured the runtime and number of operations for each approach (Fig. 5). Although the number of FLOPs reflects the theoretical operation count, the observed runtime differences are mainly determined by parallelization efficiency. Mean and max pooling have similar computational demands, while attention and transformer pooling, despite higher theoretical complexity, benefit from efficient GPU parallelization. In contrast, PH computations involve pairwise distance and filtration steps that are less parallelizable, leading to longer execution times. The ECT, although computationally intensive, is more amenable to parallel execution, resulting in smaller runtime overheads relative to its FLOPs.

**Figure 5:**
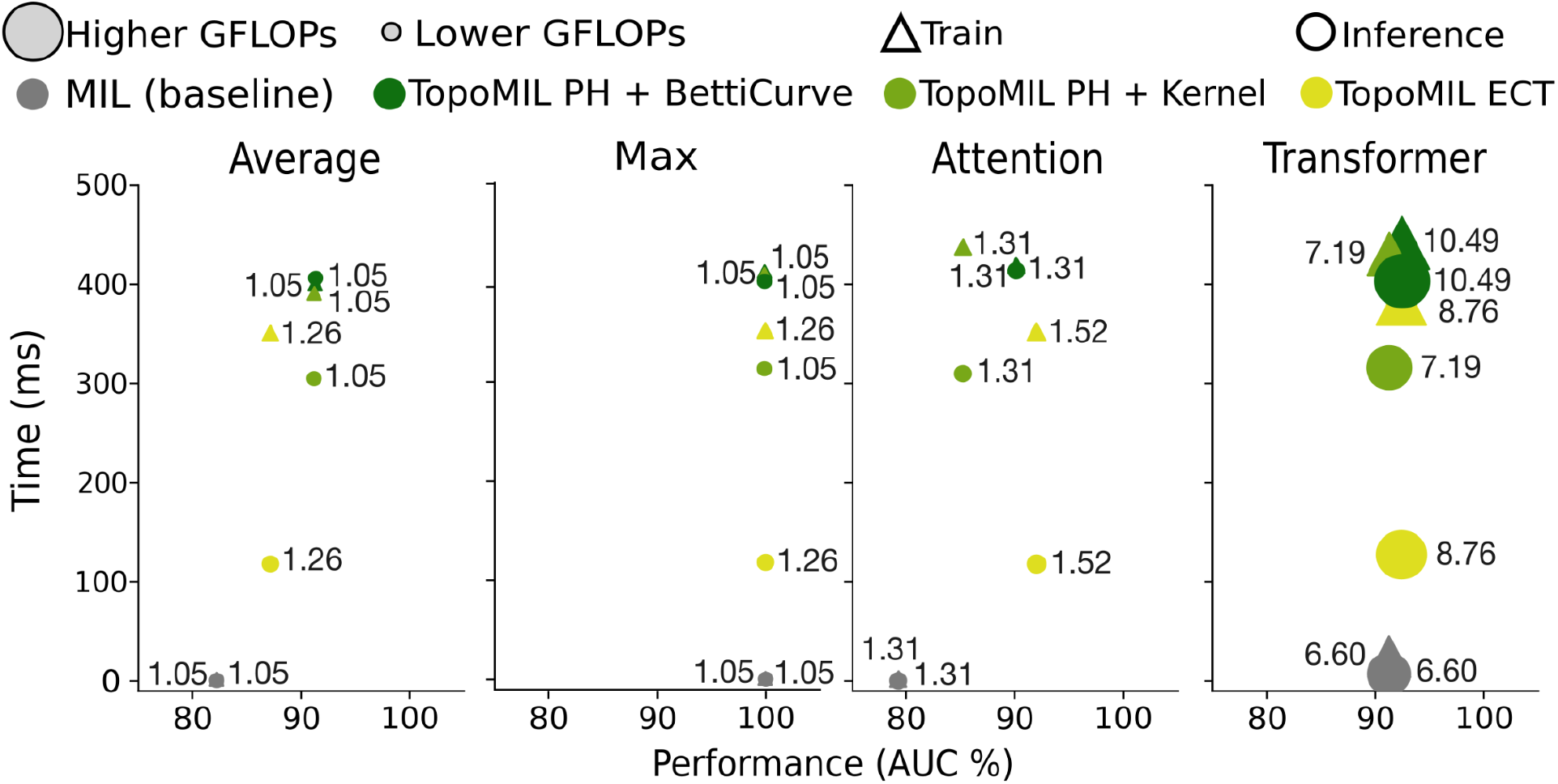
TopoMIL enhances MIL performance with trade-offs in complexity. The marker size reflects model complexity measured in GFLOPs, where larger markers indicate higher computational demands. The exact GFLOPS are written next to each shape. Different colors denote different models.

Depending on the desired balance between accuracy and efficiency, the most suitable topological method can be selected. Additional implementation details are provided in the Supplementary Information.

## Discussion

We find that bag-level topology is informative, and integrating it into multiple instance learning (MIL) improves classification performance. This aligns with the fact that the topology of instances inside a bag carries diagnostic information, which is not captured by the morphology of individual instances alone. The size and significance of its contribution however varies across the two histopathology and two cytomorphology datasets.

TopoMIL uses three approaches to compute the topological stucture of a bag. Persistent homology (PH) with a learnable kernel performs best in datasets with high heterogeneity and noise in instance representations. In contrast, the Euler characteristic transform (ECT) and PH with Betti curve yield stronger results in datasets such as Camelyon16 and Hehr, where the instance embeddings exhibit a more coherent and well-structured organization. This is supported by observations in Camelyon16, where the near-perfect performance of simple max pooling indicates that the instance encoder produces highly discriminative representations, and in Hehr, where prior analyses of the same DinoBloom foundation encoder have shown that cell embeddings cluster according to cell type, reflecting semantically consistent feature organization.

The aggregation strategy influences the effectiveness of topological descriptors. While topology captures the overall shape and relational structure of a bag, the morphological representation provides complementary information about its absolute position and orientation in the feature space, which remains critical for accurate classification. Average pooling tends to dilute class-relevant morphological features by weighting all instances equally, thereby weakening the discriminative power of the aggregated bag representation. When instance sampling is used for topology estimation, as in Camelyon16, additional variability is introduced because repeated draws from the same bag yield slightly different topologies. Combined with less discriminative morphological representations, this heterogeneity complicates classification and limits the benefit of detailed topological descriptors such as Betti curves and ECT. Nevertheless, these descriptors still provide statistically significant improvements over average pooling. The kernel-based approach, owing to its flexibility in handling heterogeneity, provides higher improvement rates under such conditions.

Attention-based aggregation partly mitigates the limitations of average pooling by emphasizing class-relevant instances, allowing the classifier to better exploit the discriminative information captured by ECT and Betti-curve descriptors. This results in stronger performance than the kernel-based method, particularly on datasets like Camelyon16 and Hehr. However, in the BRACS dataset, where classification is more challenging due to high inter-class variability and a greater dependence on holistic bag-level characteristics, the kernel-based approach remains the most effective.

Our experiments also show that transformer-based MIL achieves strong performance across most datasets by explicitly modeling interactions between instances within a bag. Integrating topological information into transformer-based MIL provides only marginal additional gains, suggesting that topology and self-attention capture related aspects of instance interactions. In contrast, incorporating topological information into simpler aggregation schemes substantially narrows or even eliminates the performance gap to transformer-based MIL, while avoiding its higher computational cost and practical constraints, such as limited sequence length.

Topology calculation is computationally complex, so integrating it into MIL requires careful consideration of computational resources. Among the three approaches, ECT is fully parallelizable and therefore faster, though it incurs higher memory usage. PH-based approaches, on the other hand, extract higher-level topological descriptors that are more robust to noise and heterogeneity, but their computational complexity remains greater.

Integrating topological information into microscopic image analysis, as presented in this work, opens new perspectives for future diagnostic applications, particularly in scenarios where global structural information is clinically important.

## Acknowledgments

The authors thank Ernst Roell for the fruitful discussions on the ECT approach. The authors also thank MLL for providing the in-house data used in our experiments.

## Author contribution

SK and BR conceived the initial idea of TopoMIL, which was further developed by SK. SK preprocessed and analyzed datasets, developed the TopoMIL framework, conducted experiments, analyzed results, wrote the manuscript, and designed the figures with input from MFD, CM, and BR. CM supervised the project, providing guidance on experimental design and manuscript structure. MFD contributed medical expertise for the cytomorphology section, performed analysis of the in-house dataset, annotated class-relevant cells for data figures, and provided the initial transformer-based code for cytomorphology data. All authors contributed to manuscript editing and have read and approved the final version.

## Methods

### TopoMIL architecture

The architecture of TopoMIL builds upon the MIL framework, where each bag consists of a collection of instance images and a label for the entire bag^29^. It comprises four main components (Fig. 1a): an instance encoder, an aggregator, a topology estimator, and a classifier head.

An **instance encoder** embeds high-dimensional instance images 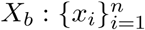 into low-dimensional representations 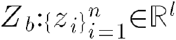.

For histopathological datasets (Camelyon16 and BRACS), we follow the CLAM^30^ baseline for preprocessing WSIs, including region-of-interest segmentation and patch extraction. For the instance embedding, we use the UNI^31^ feature extractor to embed WSI patches in a 1024-dimensional space, which has demonstrated superior performance as a foundation model (Fig. 1a).

For cytomorphology datasets, we use the DinoBloom^8^ foundation model’s feature extractors (ViT-S and ViT-B) to embed white blood cell images of peripheral blood smears (into 384-dimensional and 768-dimensional) feature spaces, respectively, for the Hehr and our in-house datasets. DinoBloom^8^ has demonstrated state-of-the-art performance when paired with a multi-attention aggregation mechanism.

**Aggregation functions** process the low-dimensional representation of instances *Z*_*b*_ ∈ ℝ ^*l*^ to generate a holistic representation of the bag *ζ*_b_ ∈ ℝ ^*ĺ*^. We consider different aggregation functions for our experiments, including average, max, attention-based, and transformer-based pooling (detailed formulations are provided in the Supplementary information).

### Topology estimation

We introduce a topology estimator block into the MIL framework, in which the instance embeddings of each bag form a point cloud in the latent space ℝ^*l*^ .This block encodes the bag’s underlying topological structure as a vector *τ*_*b*_ ∈ ℝ^*t*^ . We implement the topology estimator using two approaches based on persistent homology and a distinct approach based on the Euler characteristic transform.

### Persistent homology (PH) for topology description

This consists of two steps: (i) extraction of topological features using PH, and (ii) vector representation of the resulting persistence information.

Considering the point cloud of the bag in the latent space *Z*_*b*_ = {*z*_1_,*z*_2_, …,*z*_*n*_} the Vietoris-Rips (VR)^32^ complex at each neighborhood distance *ϵ* yields all subsets of such that the pairwise distance between subset points is less than or equal to *ϵ*.

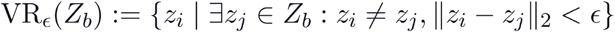

We apply the PH algorithm to the Vietoris-Rips of the input point cloud to represent its topology. PH^33^ tracks the evolution of topological features across different and provides a persistence diagram containing pairs (*ϵ*_*i*_, *ϵ*_*j*_) specifying the distance scales at which these features are created and destroyed (Fig. 1b).

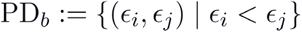

Since the number of persistent pairs *p* differs between different bags, a representation method is required to convert them into a fixed-size vector suitable for use in deep MIL models. For vectorization, we employ two approaches: (i) Betti Curve^34^ with fixed number thresholds and (ii) a learnable representation kernel^35^.

**Betti curve representation**^34^ is a statistical vectorization approach for persistence diagram ^32^. It sums the contributions of each topological feature in the persistence diagram that is active at a given *m* threshold. The Betti curve function *β* (*m*) is defined as

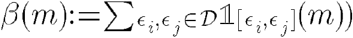

Where 𝒟 denotes the set of all birth-death pairs (*ϵ*_*i*_, *ϵ*_*j*_), and the indicator function is

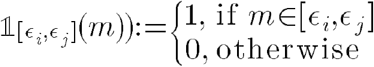

We divide the range of Betti curve values into a fixed number *t* of thresholds to ensure that all represented vectors have the same size. This is necessary because the classifier requires a fixed size of the topology vector *τ*_*b*_ (Fig. 1c).

### Learnable representation kernel

While the Betti Curve provides a simple, fixed representation, similar to other statistical methods, it is agnostic to the objective task. When dealing with deep MIL classifiers, a learnable vector representation of a persistence diagram can offer a more adaptable representation.

We define the learnable representation kernel as the probabilistic function *f* that takes *p* persistent pairs from the persistence diagram and learns the dimensional representation of that. The function is:

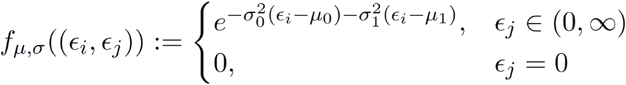

where *μ, σ* are parameters of *f*, each introducing *t* learnable mean and standard deviation for modeling the probability distribution of persistence pairs (Fig. 1d).

This is similar to the approach proposed by Hofer et al^36^, except that the noise-handling function has been removed. Since the bag’s topology is derived from the latent representations of instance images, making additional noise suppression unnecessary for the classification task.

**Euler characteristic transformation (ECT) for topology representation** proposed by Roel et. al. is a scalable ECT-based method integrable in deep learning architectures^37^. Given a point cloud of instance representations, this method scans the point cloud along multiple directions *ξ* ∈ [− *π, π*] by progressively spanning a hyperplane *h*_*ξ*_ through the space and counting the number of points it passes at each step. This produces a curve defined as

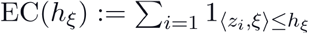

By repeating this process across multiple directions and stacking the resulting curves, a two-dimensional ECT is constructed. This topological signature is then passed through a multilayer perceptron (MLP), which compresses it into a compact *t* -dimensional representation (see Fig. 1e).

### Classifier

The two vectors, representing the bag’s morphological representation *ζ*_b_ ∈ ℝ ^*ĺ*^ and topological representation *τ*_*b*_ ∈ ℝ^*t*^, are concatenated after standardization, and the resulting *l* + *t* -dimensional vector forms the input to the classifier function.

The classification head is implemented as a two-layer multilayer perceptron (MLP) with a ReLU activation. It first projects the input vector into a 64-dimensional space, applies the ReLU non-linearity, and then maps the result to a vector of size equal to the number of output classes. The output is subsequently passed through a softmax function to obtain the predicted class probabilities.

### TopoMIL with transformer-based pooling

Due to the transformer’s capability to incorporate topology as a shape map of the entire bag, we explore an alternative design in which the bag’s topological representation is concatenated with each instance representation before being passed to the transformer. This approach enriches each instance embedding with global topological structure, enabling the self-attention mechanism to model inter-instance relationships informed not only by feature similarity but also by the overall organization of the instance space. By embedding this global-local fusion directly into the instance representations, the model gains an inductive bias that encourages learning interactions guided by the bag’s underlying topological structure. In this way 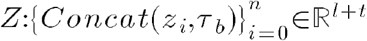 would be the input of the transformer. Then the resulting aggregated representation *ζ*_b_ ∈ ℝ ^*ĺ*^ is fed to the classification function.

### Effect of the number of topological elements

For the Betti curve, we produce fixed-length vectors by sampling a set of threshold values between the smallest and largest relevant distances within each sample. These thresholds define the curve’s resolution and representation vector size, thus serving as a key hyperparameter. Using more thresholds captures finer topological variations but can also introduce noise and increase computational cost. We empirically screened several values (100, 150, 200, 300) and found that 200 thresholds provided a good balance between stability and expressiveness for the Camelyon16 dataset, while 150 thresholds worked best for the other datasets, where higher resolutions tended to amplify dataset-specific noise due to greater heterogeneity.

In the kernel-based approach, the number of Gaussian components determines the dimensionality and granularity of the topological representation. A larger number of components captures more detailed topological variations, while fewer components yield smoother representations that are more robust to noise. Since heterogeneous datasets are more prone to noise in detailed topological representations, using fewer Gaussian components helps produce more stable and reliable embeddings. We empirically evaluated different numbers of Gaussian components (20, 40, 60, 100, 150) and found that 20 components performed best for BRACS, which exhibits higher variability, while 60 components provided the best trade-off between detail and stability for the other datasets.

For the ECT, three hyperparameters are introduced: the number of orientations used to scan the latent space, the sharpness factor of the sigmoid function, and the dimensionality of the output vector. We set the number of orientations equal to the embedding space dimensionality, use a sharpness factor of 256, and define the representation vector to have 100 dimensions. These values were determined heuristically based on preliminary experiments showing stable and consistent performance across datasets.

## Code availability

We publicly release the TopoMIL framework on GitHub at https://anonymous.4open.science/r/TopoMIL_final-6BCB/.

## References

1. Elhassan, T. A. M., Mohd Rahim, M. S., Osman, A. H., Chaudhri, N. A. & Aljurf, M. A deep learning framework for classification of immature white blood cells in acute myeloid leukemia using convolutional Autoencoder and transfer-learning ensemble. Blood 142, 7290–7290 (2023).

2. Park, S., Park, Y. H., Huh, J., Baik, S. M. & Park, D. J. Deep learning model for differentiating acute myeloid and lymphoblastic leukemia in peripheral blood cell images via myeloblast and lymphoblast classification. Digit. Health 10, 20552076241258079 (2024).

3. He, L., Long, L. R., Antani, S. & Thoma, G. R. Histology image analysis for carcinoma detection and grading. Comput. Methods Programs Biomed. 107, 538–556 (2012).

4. Tian, F. et al. Prediction of tumor origin in cancers of unknown primary origin with cytology-based deep learning. Nat. Med. 30, 1309–1319 (2024).

5. Carbonneau, M.-A., Cheplygina, V., Granger, E. & Gagnon, G. Multiple instance learning: A survey of problem characteristics and applications. Pattern Recognit. 77, 329–353 (2018).

6. Waqas, M., Ahmed, S. U., Tahir, M. A., Wu, J. & Qureshi, R. Exploring Multiple Instance Learning (MIL): A brief survey. Expert Syst. Appl. 250, 123893 (2024).

7. Cisternino, F. et al. Self-supervised learning for characterising histomorphological diversity and spatial RNA expression prediction across 23 human tissue types. Nat. Commun. 15, 5906 (2024).

8. Koch, V. et al. DinoBloom: a foundation model for generalizable cell embeddings in hematology. in International Conference on Medical Image Computing and Computer-Assisted Intervention 520–530 (Springer, 2024).

9. Ilse, M., Tomczak, J. & Welling, M. Attention-based deep multiple instance learning. ICML 80, 2132–2141 (2018).

10. Shao, Z. et al. TransMIL: Transformer based correlated multiple instance learning for whole slide image classication. Neural Inf Process Syst 34, 2136–2147 (2021).

11. Gardić, N. G., Miljković, D. M. & Lovrenski, A. N. Cytomorphological features as a subtyping tool of non-small-cell lung cancer in brushing bronchoscopic samples. J. Cytol. 41, 143–149 (2024).

12. Lin, S. et al. Digital quantification of tumor cellularity as a novel prognostic feature of non-small cell lung carcinoma. Mod. Pathol. 36, 100055 (2023).

13. Bukkuri, A., Andor, N. & Darcy, I. K. Applications of topological data analysis in oncology. Front. Artif. Intell. 4, 659037 (2021).

14. Estey, E., Hasserjian, R. P. & Döhner, H. Distinguishing AML from MDS: a fixed blast percentage may no longer be optimal. Blood 139, 323–332 (2022).

15. Chulián, S. et al. The shape of cancer relapse: Topological data analysis predicts recurrence in paediatric acute lymphoblastic leukaemia. PLoS Comput. Biol. 19, e1011329 (2023).

16. Wang, Q., Song, A., Batsivari, A., Bonnet, D. & Monod, A. A topological Gaussian mixture model for bone marrow morphology in leukaemia. arXiv [stat.AP] (2025).

17. Matek, C., Schwarz, S., Spiekermann, K. & Marr, C. Human-level recognition of blast cells in acute myeloid leukaemia with convolutional neural networks. Nat Mach Intell 1, 538–544 (2019).

18. Kömen, J., Marienwald, H., Dippel, J. & Hense, J. Do histopathological foundation models eliminate batch effects? A comparative study. arXiv [cs.LG] (2024).

19. Wasserman, L. Topological Data Analysis. arXiv [stat.ME] (2016) doi:10.1146/annurev-statistics-031017-100045.

20. Chazal, F., de Silva, V. & Oudot, S. Persistence stability for geometric complexes. Geom. Dedicata 173, 193–214 (2014).

21. Chazal, F., Cohen-Steiner, D., Guibas, L. J., Mémoli, F. & Oudot, S. Y. Gromov-Hausdorff Stable Signatures for Shapes using Persistence. Comput. Graph. Forum 28, 1393–1403 (2009).

22. Obeid, A. et al. PMIL: A topology module to improve MIL-based WSI classification. in 2025 IEEE International Symposium on Circuits and Systems (ISCAS) 1–5 (IEEE, 2025). doi:10.1109/iscas56072.2025.11043738.

23. Koung, P. F., Fatema, S., Pakasticali, N., Luu, H. S. & Coskunuzer, B. Diagnosis of blood diseases and disorders with topological deep learning. medRxiv (2025) doi:10.1101/2025.01.21.25320908.

24. Kazeminia, S., Marr, C. & Rieck, B. Topologically regularized Multiple Instance Learning to harness data scarcity. arXiv [cs.LG] (2023).

25. Bejnordi, B. E. et al. Diagnostic assessment of deep learning algorithms for detection of lymph node metastases in women with breast cancer. JAMA 318, 2199–2210 (2017).

26. Brancati, N. et al. BRACS: A dataset for BReAst Carcinoma Subtyping in H&E histology images. Database: The Journal of Biological Databases and Curation 2022, (2021).

27. Hehr, M. et al. Explainable AI identifies diagnostic cells of genetic AML subtypes. PLOS Digit. Health 2, e0000187 (2023).

28. Dasdelen, M. F. et al. Transformer-based hematological malignancy prediction from peripheral blood smears in a real-world cohort. arXiv [q-bio.QM] (2025).

29. Dietterich, T. G., Lathrop, R. H. & Lozano-Pérez, T. Solving the multiple instance problem with axis-parallel rectangles. Artif. Intell. 89, 31–71 (1997).

30. Lu, M. Y. et al. Data-efficient and weakly supervised computational pathology on whole-slide images. Nat. Biomed. Eng. 5, 555–570 (2021).

31. Chen, R. J. et al. Towards a general-purpose foundation model for computational pathology. Nat. Med. 30, 850–862 (2024).

32. Hensel, F., Moor, M. & Rieck, B. A survey of topological machine learning methods. Front. Artif. Intell. 4, 681108 (2021).

33. Barannikov, S. The framed Morse complex and its invariants. Advances in Soviet Mathematics 21, 93–116 (1994).

34. Chevyrev, I., Nanda, V. & Oberhauser, H. Persistence paths and signature features in topological data analysis. IEEE Trans. Pattern Anal. Mach. Intell. 42, 192–202 (2020).

35. Hofer, C., Kwitt, R., Niethammer, M. & Uhl, A. Deep learning with topological signatures. Neural Inf Process Syst abs/1707.04041, (2017).

36. Hofer, C., Kwitt, R., Niethammer, M. & Uhl, A. Deep learning with topological signatures. Advances in neural information processing systems 30, (2017).

37. Roell, E. & Rieck, B. Differentiable Euler characteristic transforms for shape classification. arXiv [cs.LG] (2023).

38. Wagner, S. J. et al. Transformer-based biomarker prediction from colorectal cancer histology: A large-scale multicentric study. Cancer Cell 41, 1650–1661.e4 (2023).

