## Supplementary information for "TopoMIL: Topology Improves Multiple Instance Learning in Diagnostic Microscopic Images"

### Datasets

The **Camelyon16**[25] dataset contains 400 hematoxylin and eosin (H&E)-stained WSIs of lymph node sections collected for the detection of breast cancer metastases (Fig. 2a). It contains 159 slides with tumor regions, and 241 slides of normal tissue. Each WSI contains about 1,000 to 100,000 patches. The **BRACS**[26] dataset includes 547 H&E-stained WSIs of breast tissue lesions taken from 189 patients, annotated into three diagnostic categories: 265 benign, 193 malignant, and 89 atypical cases (Fig. 2b). The slides exhibit varying tissue sizes and staining characteristics, resulting in bags containing 500 to 25,000 patches. Each WSI is divided into patches serving as instances, where each patch may contain cells or other histological components, depending on the sampled region. Since adjacent patches often share contextual information, instances within a WSI exhibit spatial dependence.

Each cytomorphology dataset comprises microscopic images of blood samples, with each sample treated as a bag. Instances correspond to segments centered on individual white blood cells, often accompanied by red blood cells. In this setting, instances are spatially independent of one another. The **Hehr**[27] dataset is derived from peripheral blood smears of patients diagnosed with acute myeloid leukemia (AML), labeled according to their genetic subtype, as well as from healthy controls. The dataset includes 189 patient-level samples across five categories: 30 cases with CBFB–MYH11 fusion, 40 cases of NPM1-mutated AML, 30 cases of acute promyelocytic leukemia (APL; PML–RARA fusion), 29 cases of core-binding factor AML with RUNX1–RUNX1T1 fusion, and 60 samples from healthy individuals. The number of WBC images per sample ranges from 99 to 500 WBC images per bag, with a mean of 430±107 cells. Our **in-house**[28] dataset comprises peripheral white blood cell images collected from 2043 patients diagnosed across eight broad hematologic categories: acute leukemias (AL), myelodysplastic syndromes (MDS), overlapping MDS/myeloproliferative neoplasms (MDS/MPN), myeloproliferative neoplasms (MPN), lymphoma, plasma cell neoplasms (PCN), reactive changes, and normal cases from stem cell donors. This dataset contains a larger cohort of cytomorphology images. However, it also introduces a higher level of semantic ambiguity due to the larger class overlaps. We split the dataset into training and test sets using an 80:20 ratio, stratified by diagnostic group (409 patients reserved for testing). The number of WBC images per patient ranges from 55 to 500, with a mean of 488±55 cells.

### Data availability

Three of four datasets used in this study are publicly available. For histopathology experiments, we used Camelyon16 for lymph node metastasis detection. We followed preprocessing steps similar to those used in the CLAM framework, including patch extraction from whole-slide images; further details are available at the CLAM GitHub repository. Additionally, we used the BRACS dataset for breast cancer subtyping, accessible at BRACS website. For cytomorphology, we employed the Hehr dataset, published alongside the SCEMILA framework by Hehr et al., available at GitHub. Our in-house cytomorphology dataset is not publicly available due to privacy restrictions.

### Aggregation functions

- **Average pooling** calculates the bag's morphological representation by averaging the features of all its instances of the bag:

  $\zeta_b := \frac{1}{n}\sum_{i=1}^{n} z_i$

- **Max pooling** calculates the bag's morphological representation as the element-wise maximum feature present in its instances:

  $\zeta_b := \max_{1\leq i\leq n} z_i$

- **Attention-based pooling** calculates the bag's morphological representation by a weighted averaging over its instances as:

  $\zeta_b := \sum_{i=1}^{n} a_i.z_i,$

  where $a_i$ is the attention value for the $i$th instance of the bag, calculated as

  $a_i := \frac{\exp(W_1.tanh(W_2.z_i))}{\sum_{j=1}^{n}\exp(W_1.tanh(W_2.z_j))}.$

  where $W_1$ and $W_2$ are learnable parameters of the attention network. We used this approach for histopathology datasets, while for the cytomorphology datasets, we use multi-attention functions as they provide state-of-the-art performance[8] on these data. This function calculates the attention value $a_{i,c}$ for instance $z_i$ and class $c \in C$:

  $\zeta_b^c := \sum_{i=1}^{n} a_{i,c}.z_i,$

  where $a_i^c$ is calculated as:

  $a_{i,c} := \frac{\exp(W_1^c.tanh(W_2^c.z_i))}{\sum_{j=1}^{n}\exp(W_1^c.tanh(W_2^c.z_j))}$

  where and $W_1^c$ and $W_2^c$ are learnable attention network parameters for each class $c \in C$.

- **Transformer-based pooling** employs the concept of the self-attention mechanism that captures the relative dependencies among all instances within a bag. This mechanism introduces the concepts of quarry ($Q$), key ($K$), and value ($V$), each computed as a learned linear transformation of instance embeddings via weight matrices $W_Q \in \mathbb{R}^{l\times l_k}$, $W_K \in \mathbb{R}^{l\times l_k}$, and $W_V \in \mathbb{R}^{l\times l_v}$. Given a sequence of instance embeddings $Z = \{z_i\}_{i=1}^{n}$ with $z_i \in \mathbb{R}^{l}$, the transformations are defined as:

  $Q = W_Q.Z$, $K = W_K.Z$, $V = W_V.Z$,

  The self-attention operation is defined as:

  $\hat{S}(Q,K,V) = softmax(\frac{Q.K^T}{\sqrt{l_k}}).V$,

  which aggregates the values based on the similarity between queries and keys, thus capturing instance correlations within the embedding space.

For the cytomorphology dataset, we use a recently published transformer-based MIL, which applies a multi-head self-attention as:

$$\zeta_b = Concat_{h=1}^{H}(\hat{S}(Q_h, K_h, V_h)).W_o,$$

where $w_o \in \mathbb{R}^{H.l_v \times l}$ and $H$ denote the number of self-attention heads working in parallel. Similar to many other approaches, two transformer layers, each containing 8 self-attention heads, are employed. Positional embeddings are omitted in our setting due to the spatial independence of instances within blood samples.

For Histopathology datasets, we use the transMIL[10] architecture, which incorporates a pyramid positional encoding generator to model spatial dependencies among instances. This approach also introduces a learnable classification token $z_c \in \mathbb{R}^l$ prepended to the sequence of instance embeddings $Z = [z_c, z_1, .., z_n]$. After the first transformation layer, the spatial information of instances is fused before the sequence is processed by a second transformer layer for final aggregation.

### Evaluation metrics

For evaluation, we used the ROAUC (Receiver Operating Characteristic Area Under the Curve) metric, which measures a model's ability to distinguish between positive and negative classes across all possible classification thresholds. It is computed by plotting the true positive rate against the false positive rate at different thresholds and calculating the area under the resulting curve. A higher ROAUC indicates better discriminative performance, with 1.0 representing perfect separation and 0.5 representing random guessing. Its threshold-independent nature makes ROAUC particularly suitable for models, like MIL, that produce continuous prediction scores.

**Statistical analysis**. To assess the statistical significance of TopoMIL improvement across multiple datasets and aggregations, we used the Kruskal-Wallis test, a non-parametric method for comparing more than two independent samples. It does not assume normality, making it suitable for metrics such as ROAUC that may not follow a Gaussian distribution. The test proceeds as follows: all observations from all groups are ranked in ascending order; tied values are assigned the average of their ranks. For each group, the sum of ranks $R_i$ and the number of observations $n_i$ are computed. The Kruskal--Wallis statistic $H$ is then calculated as:

$$H = \frac{12}{N(N+1)} \sum_{i=1}^{k} \frac{R_i^2}{n_i} - 3(N+1)$$

where $N$ is the total number of observations across all groups and $k$ is the number of groups. Under the null hypothesis that all groups come from the same distribution, $H$ approximately follows a chi-squared distribution with $k-1$ degrees of freedom. A significant p-value (<0.05) indicates that at least one group's performance differs from the others.

| Dataset | Method | Average pooling | Max pooling | Attention-based pooling | Transformer-based pooling |
|---|---|---|---|---|---|
| camelyon | MIL | 83.0 ± 6.6 | 97.7 ± 1.0 | 78.2 ± 5.3 | 92.1 ± 4.9 |
| | TopoMIL PH+BettiCurve | 91.5 ± 4.4 | 99.1 ± 0.4 | 90.4 ± 5.1 | **92.7 ± 5.5** |
| | TopoMIL PH+Kernel | **91.1 ± 2.8** | 99.7 ± 0.3 | 85.2 ± 4.9 | 91.3 ± 4.7 |
| | TopoMIL ECT | 87.9 ± 4.6 | **99.7 ± 0.2** | **91.3 ± 3.8** | **92.7 ± 5.5** |
| BRACS | MIL | 78.3 ± 0.4 | 76.8 ± 1.6 | 71.4 ± 2.5 | 72.5 ± 2.3 |
| | TopoMIL PH+BettiCurve | 78.1 ± 1.0 | 76.7 ± 1.2 | 70.8 ± 2.9 | 73.0 ± 3.0 |
| | TopoMIL PH+Kernel | **79.2 ± 0.5** | **77.3 ± 0.7** | **78.4 ± 1.3** | **73.4 ± 1.8** |
| | TopoMIL ECT | 77.9 ± 0.3 | 77.9 ± 1.4 | 70.7 ± 1.5 | 73.7 ± 3.5 |
| Hehr | MIL | 93.1 ± 0.6 | 75.4 ± 1.6 | 93.8 ± 1.6 | 97.3 ± 1.2 |
| | TopoMIL PH+BettiCurve | **95.4 ± 0.4** | 76.3 ± 3.5 | **96.7 ± 0.7** | 96.6 ± 0.8 |
| | TopoMIL PH+Kernel | 95.3 ± 0.7 | 74.5 ± 2.1 | **96.6 ± 0.5** | 97.3 ± 1.0 |
| | TopoMIL ECT | 95.0 ± 0.5 | **86.6 ± 1.0** | 96.5 ± 0.6 | **97.3 ± 0.4** |
| In-house | MIL | 87.2 ± 1.3 | 82.4 ± 1.0 | 86.2 ± 2.0 | 89.3 ± 1.1 |
| | TopoMIL PH+BettiCurve | 88.1 ± 1.0 | **84.9 ± 1.3** | 86.7 ± 1.2 | 89.4 ± 1.9 |
| | TopoMIL PH+Kernel | 87.8 ± 0.5 | 84.7 ± 1.7 | 86.8 ± 0.9 | **89.4 ± 1.3** |
| | TopoMIL ECT | **88.6 ± 1.2** | 84.7 ± 1.0 | **86.8 ± 0.9** | 88.8 ± 1.1 |

**Table 1: TopoMIL enhances MIL performance across diverse biomedical imaging datasets and aggregation functions.** Values represent the mean ± standard deviation of AUROC (%), with the best performance for each dataset and aggregation function highlighted in bold.

| Dataset | Method TopoMIL | Average pooling | Max pooling | Attention_based pooling | Transformer-based pooling |
|---|---|---|---|---|---|
| camelyon | PH+BettiCurve | **0.0041** | 0.5032 | **0.0005** | 0.5202 |
| | PH+Kernel | **0.0011** | 0.4656 | **0.02818** | 0.7051 |
| | ECT | 0.0755 | 0.9421 | **0.0002** | 0.6501 |
| BRACS | PH+BettiCurve | **0.0040** | 0.5032 | **0.0005** | 0.5202 |
| | PH+Kernel | **0.0011** | 0.4656 | **0.02819** | 0.7051 |
| | ECT | 0.0755 | 0.9422 | **0.0002** | 0.6501 |
| Hehr | PH+BettiCurve | **0.0090** | 0.3472 | **0.0163** | 0.2506 |
| | PH+Kernel | **0.0162** | 0.6015 | **0.0163** | 0.7540 |
| | ECT | **0.0163** | **0.0090** | **0.0090** | 0.9168 |
| In-house | PH+BettiCurve | 0.3472 | **0.0090** | 0.4647 | 0.4647 |
| | PH+Kernel | 0.4647 | **0.0283** | 0.3472 | 0.9168 |
| | ECT | 0.1172 | **0.0090** | 0.4647 | 0.2506 |

**Table 2: Significance of the enhancement made by TopoMIL**. p-values are calculated based on the Kruskal-Wallis test. Significant enhancements are bolded.

| Dataset | MIL | TopoMIL PH+BettiCurve | TopoMIL PH+Kernel | TopoMIL ECT |
|---|---|---|---|---|
| Camelyon | **86.8 ± 8.7** | 61.4 ± 7.7 | 76.2 ± 8.9 | 74.5 ± 6.7 |
| BRACS | **74.6 ± 3.7** | 45.4 ± 5.2 | 56.4 ± 7.2 | 65.1 ± 2.3 |
| Hehr | 85.3 ± 9.6 | 55.0 ± 7.6 | 76.6 ± 4.4 | **91.3 ± 3.5** |
| In-house | **85.8 ± 3.1** | 51.5 ± 3.0 | 71.9 ± 1.0 | 85.0 ± 1.7 |

**Table 3: Ablation study,** evaluating the standalone contribution of topological representations across all datasets and approaches (BettiCurve, kernel, and ECT), showing that ECT provides the strongest signal, kernel yields moderate improvements, and BettiCurve contributes the least. Performances are reported as mean ± std of AUROC (%).

### Computational complexity and scalability

We analyze the computational cost of the proposed framework through both theoretical analysis and empirical evaluation (Fig. 5), comparing it against the baseline MIL framework. In this study, all reported results are based on 0-dimensional PH features, whose computational complexity is dominated by the pairwise distance computations between $n$ instances in a bag represented in a $d$-dimensional space, yielding a time complexity of $\mathcal{O}(n^2.d)$. In our implementation, parallelization reduces the effective time complexity to $\mathcal{O}(n.d)$, while the memory complexity remains $\mathcal{O}(n^2)$ for storing the distance matrix. The Betti curve computation adds only a constant-factor overhead, while the learnable kernel introduces an $\mathcal{O}(e)$ term, with $e$ as the number of probabilistic elements ($20 < e < 200$), which is dominated by the complexity of PH.

For the ECT approach, given a bag with $n$ points in a latent space of d-dimensional latent space, the main cost comes from projecting the point cloud onto $d$ uniformly sampled directions and evaluating the transform over $d$ resolution bins. This yields an overall time complexity of $\mathcal{O}(n.d^2)$, which is linear in the number of points but quadratic in the latent dimensionality. The memory complexity of $\mathcal{O}(n + d^2)$ for storing the point cloud and the transform values.

This relatively high computational cost motivated us to use a random subset of 1000 instances per bag for the Camelyon16 dataset, while full-bag computation is feasible for other three smaller datasets.

Max-pooling and average-pooling require $\mathcal{O}(n.d)$ time and minimal additional memory. Attention-based pooling computes weights for all $n$ instances, resulting in $\mathcal{O}(n.d + n)$ time and $\mathcal{O}(n)$ memory for storing the weights. Transformer-based pooling involves pairwise interactions between all instances, with time complexity $\mathcal{O}(n^2.d)$ and memory complexity $\mathcal{O}(n^2)$ for the attention matrix. While simple pooling operations scale linearly with bag size, transformer-based pooling scales quadratically, making it computationally demanding for large bags and often necessitating instance subsampling or hierarchical strategies. Due to the high computational cost of transformer-based aggregation, integrating topological estimation on top of it requires applying our framework on a (1%) subsample of instances per bag. This subsampling ensures that both morphological and topological embeddings of bags remain computationally feasible.

All experiments were conducted on an HPC cluster using NVIDIA A100 and H100 GPUs. For the Camelyon16 dataset, we employed distributed training across four GPUs to handle its large data volume, resulting in an effective batch size of four. All evaluations were performed on a single GPU with a batch size of one.
